# Tactile Adaptation to Orientation Produces a Robust Tilt Aftereffect and Exhibits Crossmodal Transfer When Tested in Vision

**DOI:** 10.1101/2023.12.20.572503

**Authors:** Guandong Wang, David Alais

## Abstract

Orientation processing is one of the most fundamental functions in both visual and somatosensory perception. Converging findings suggest that orientation processing in both modalities is closely linked: somatosensory neurons share a similar orientation organisation as visual neurons, and the visual cortex has been found to be heavily involved in tactile orientation perception. The tilt aftereffect (TAE) is a demonstration of orientation adaptation and is used widely in behavioural experiments to investigate orientation mechanisms in vision. By testing the classic TAE paradigm in both tactile and crossmodal orientation tasks between vision and touch we were able to show that tactile perception of orientation shows a very robust TAE, similar to its visual counterpart. We further show that orientation adaptation in touch transfers to produce a TAE when tested in vision, but not vice versa. We also observed a similar asymmetricity in the intramodal serial dependence effect within the test sequence. These findings provide concrete evidence that vision and touch engage a similar orientation processing mechanism, but the asymmetry in the crossmodal transfer of TAE and serial dependence provides more insights into the underlying mechanism of this link.

## Introduction

Orientation is considered one of the most basic features in vision^1^. It is extracted very early in visual processing by specialized neurons in the primary visual cortex^2^ and plays a vital role as one of the essential building blocks of visual perception. The receptive fields of neurons in the primary visual cortex (area V1) are narrowly tuned to the orientation of contours at a particular spatial scale. Collectively, the population of V1 neurons encodes all orientations over a wide range of spatial scales^3^. Similar to vision, neurons in the early somatosensory cortex (areas S1 and S2) show an orientation selectivity. Although not as specialized or narrowly tuned for orientation as V1 neurons, the population of all orientation-tuned somatosensory neurons covers all orientations at a range of scales^4–6^. Given these similarities and considering the value of efficient sensory coding, it is natural to wonder whether there is a common processing of orientation between the visual and somatosensory modalities.

Various findings have accumulated in support of the view that orientation processing in visual and somatosensory modalities is linked. First, a number of studies using various brain imaging methods have reported occipital lobe (visual cortex) involvement during purely tactile tasks with both sighted and visual-impaired individuals^7–15^. These findings all point to the involvement of extrastriate visual cortex during the processing of a variety of tactile tasks. More importantly, they reveal that the spatial scale of the task feature regulates the degree of visual involvement: large features such as size, shape and orientation (macrospatial tactile scale) tend to activate the visual cortex whereas fine-grained or microspatial tasks (e.g., texture and roughness, for which somatosensory neurons are specialised) do not. One interpretation is that macrospatial tactile information is passed onto the visual cortex because vision is more specialized for such information; and that this occurs in a top-down manner, possibly as visual imagery^7,9,13,15–17^.

Several studies have specifically tested for tactile input to the extrastriate visual cortex during orientation discrimination (macrospatial task) versus spatial frequency discrimination (microspatial task) of tactile gratings. Converging evidence from both PET^16^ and fMRI studies^15^ indicates that orientation discrimination produces a significant task-specific activation in extrastriate visual cortex compared with spatial frequency discrimination. Moreover, when transcranial magnetic stimulation (TMS) was applied to this extrastriate visual area, the participant’s ability to discriminate the tactile grating’s orientation was severely affected, but not their ability to discriminate its spatial frequency^13^. Complementing these results, a perceptual learning study of grating orientation showed tactile-to-visual transfer of orientation learning^18^. Together, these findings support the hypothesis that tactile orientation information is passed on to the extrastriate visual cortex, presumably to benefit from vision’s finer orientation acuity.

There is another line of evidence implicating connections between the somatosensory cortex and early visual cortex, including area V1. Neuronal recordings in rats have found primary somatosensory cortex (S1) and primary visual cortex (V1) carry similar amounts of information during purely tactile tasks, and V1 responses correlate strongly with task performance^19^. In humans, one study found activation in V1 and deactivation in extrastriate visual cortex during a tactile dot spacing task^20^. Complementing this, psychophysical studies show that visual contrast sensitivity for grating stimuli is improved when paired with a tactile grating that is congruent with the visual stimulus. Critically, varying the relative orientation between the gratings reveals the crossmodal facilitation has a very tight orientation tuning, implying an interaction at the earliest cortical stage where orientation tunings are tightest^21^. Another study found a similarly tight orientation tuning between touch and vision in a visual suppression paradigm in which visual gratings were less suppressed when paired with a congruent or near-congruent tactile grating^22^. These findings support the possibility of direct connections between primary visual and somatosensory cortices, such as those documented in rodent brain^23,24^, and hence an early integration of visual and tactile orientation information. Thus, while there is a considerable accumulation of evidence documenting visual-tactile interactions for orientation tasks, it is not clear at which cortical stage these interactions occur.

Adaptation and aftereffects are often used in the exploration of underlying neural mechanisms across various modalities, providing insights into phenomena such as crossmodal interaction^25–28^. In vision, orientation is often studied using the tilt aftereffect (TAE). The TAE is an old and well-known aftereffect that has revealed much about visual orientation mechanisms^29^. The TAE occurs when a tilted stimulus presented for a prolonged period causes nearby orientations to appear repelled away from the adaptor’s orientation. These perceptual repulsion effects are linked to corresponding changes in the cortical responses of V1 neurons^30,31^. These adaptation-induced neural changes, similar to many other visual aftereffects^32^, are traditionally explained by a simple fatigue model in which neurons preferring the adaptor orientation show reduced responsivity after adaptation, increasing the relative response strength to nearby orientations. Although usually studied as a visual aftereffect, a tactile TAE has been demonstrated on the palm of the hand using a two-point stimulation method^33^, and visual adaptation to tilted gratings biases perception of subsequent two-point tactile orientation stimuli^34^. In addition, combining tactile and visual surround gratings is found to increase the visual tilt illusion^35^. To date, there has been no systematic demonstration of tactile and crossmodal orientation aftereffects within the same paradigm.

In this study, we will test the TAE in touch, as well as testing for transfer between modalities to further test the proposed common orientation mechanisms in touch and vision. Our results reveal a robust tactile TAE and also show that adaptation effects can transfer between modalities in an asymmetrical way, where tactile adaptors bias subsequent visual perception, but not vice versa. This provides concrete evidence of touch and vision engaging common orientation mechanisms, while the asymmetricity of adaptation transfer is informative about the links underlying visual-tactile orientation interactions. In addition to the main findings, we also took a closer look at how the adaptation effect changes after the adaptation phase by further examination of the trial-by-trial variability during the test phase. We found a similar asymmetrical intramodal serial dependence effect^36,37^ between the test trials, which provides additional insights into the potential source of this asymmetricity.

## Results

Two experiments were conducted to assess the tactile tilt aftereffect (TAE) and the transfer of TAE between vision and touch. Figure 1 illustrates the experimental setup and example stimuli. Participants underwent an adaptation phase in which tilted stimuli (Experiment 1: tactile; Experiment 2, Condition 1: tactile, Condition 2: vision) were presented for a prolonged duration (Experiment 1: 30 s, Experiment 2: 60 s). Following this, test stimuli were presented either in the same modality (Experiment 1: tactile) or different modalities (Experiment 2, Condition 1: visual; Condition 2: touch). Participants then indicated their responses by judging the orientation of the stimulus with respect to the vertical axis (proximal-distal direction in touch). Psychometric functions were fitted for each participant under each adaptor orientation (left vs. right). The point of subjective equality (PSE) was obtained from the fitted psychometric functions and was used as the measure of adaptation-induced orientation bias, with positive values indicating a leftward bias and negative values indicating a rightward bias. The TAE is then quantified by the difference in PSE (Δ*PSE* = *PSE*_*right*_ − *PSE*_*left*_) between the two adaptor orientations (left vs. right), where positive values indicate a repulsive TAE and negative values indicate an attractive shift. See Methods for more details.

**Figure 1.**
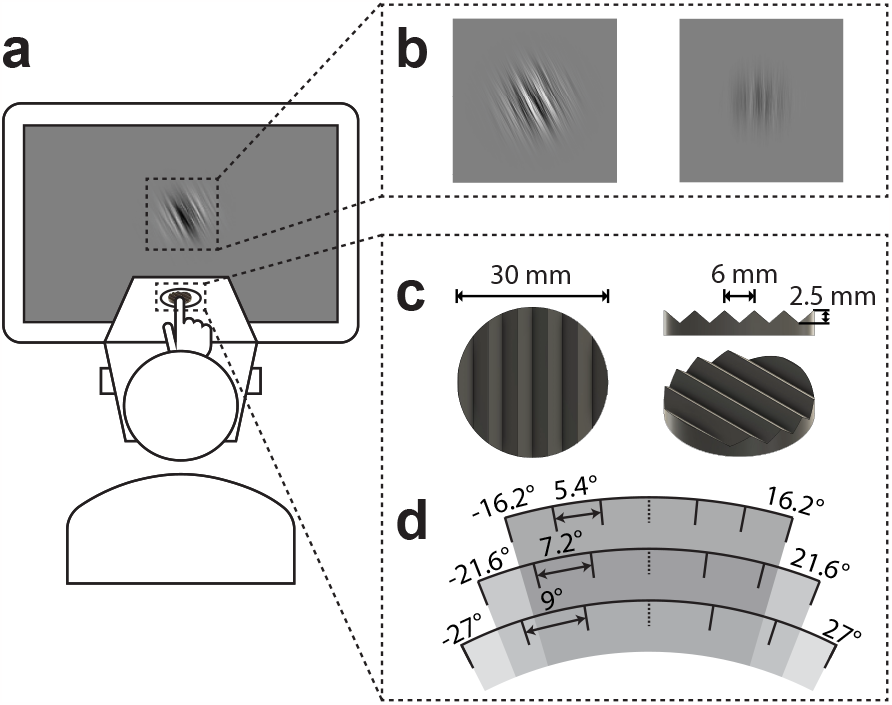
**(a)** Experimental setup. The tactile stimuli were presented via a device positioned in front of the screen and the instructions and visual stimuli were presented on the screen. **(b)** Examples of adaptation (left image) and test (right image) visual stimulus used in Experiment 2. Both types of stimulus were orientation-filtered pink noise with different RMS contrast levels (adaptation: 0.300, test: 0.018). **(c)** Illustration of the tactile grating used in both experiments. **(d)** The three sets of stimulus orientations used in Experiment 1. The sets have the same mean but vary in range. One set was used per participant, with the choice based on their performance in a preliminary test of the participant’s tactile orientation resolution. In Experiment 2, only the 3rd set (−27° to 27° with 9° gap) was used.

### Robust repulsive tilt aftereffect (TAE) in touch

Experiment 1 investigates whether adaptation to somatosensory orientation will produce a TAE, similar to what is observed in the visual system. The experiment consists of two phases: an adaptation and a test phase (Fig. 2). During the adaptation phase, participants were instructed to passively feel the orientation of a tilted tactile grating(−27°or 27°) without making any response. After adaptation, a test stimulus was presented (a rotated version of the same tactile grating), and participants indicated whether the test grating was oriented left or right with respect to the proximal-distal direction.

**Figure 2.**
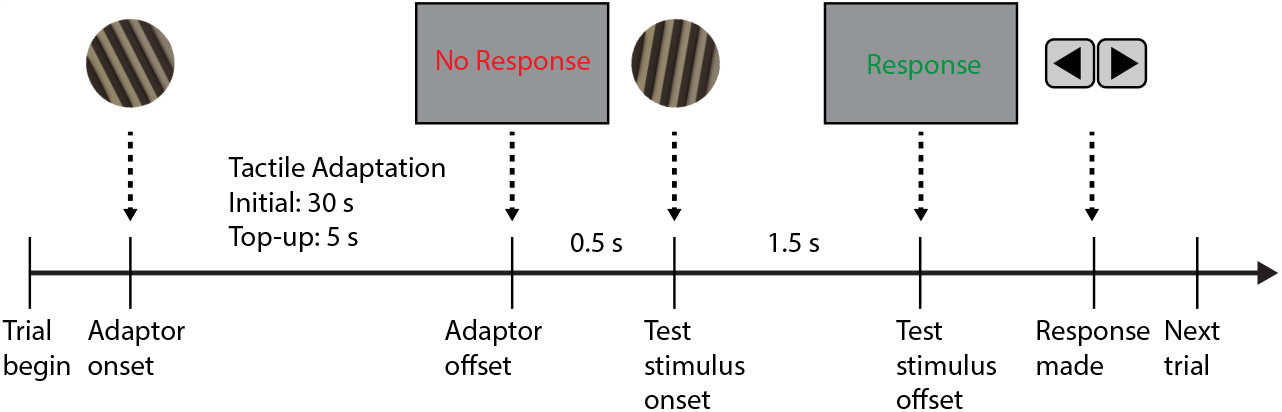
Procedures for Experiment 1, Tactile TAE: The experiment consists of two phases, an adaptation phase (initial:30 s, top-up: 5 s) and a test phase. Participants were adapted by passively feeling the orientation of a tilted grating (−27°or 27°). During the test phase, one of the 6 test stimulus orientations was presented for 1.5 s. Participants were then cued to respond using the arrow keys on the keyboard to indicate whether the perceived orientation was left or right of the vertical (proximal-distal) axis.

A permutation test on the PSE difference between leftward and rightward adaptor conditions (Fig. 3) revealed a robust repulsive perceptual shift from the adaptor orientation (Mean Δ*PSE* = 11.11°, *SD* = 5.70, *t*_*obs*_(15) = 7.30, *p*_*obs*_ *<* .001, Cohen’s *d*_*obs*_ = 1.83, *p*_*perm*_ *<* .0001^***^). The tactile TAE was strong and reliable and was present at the individual level for all 14 participants (Fig. 3a, grey dots and lines represent individual participants).

**Figure 3.**
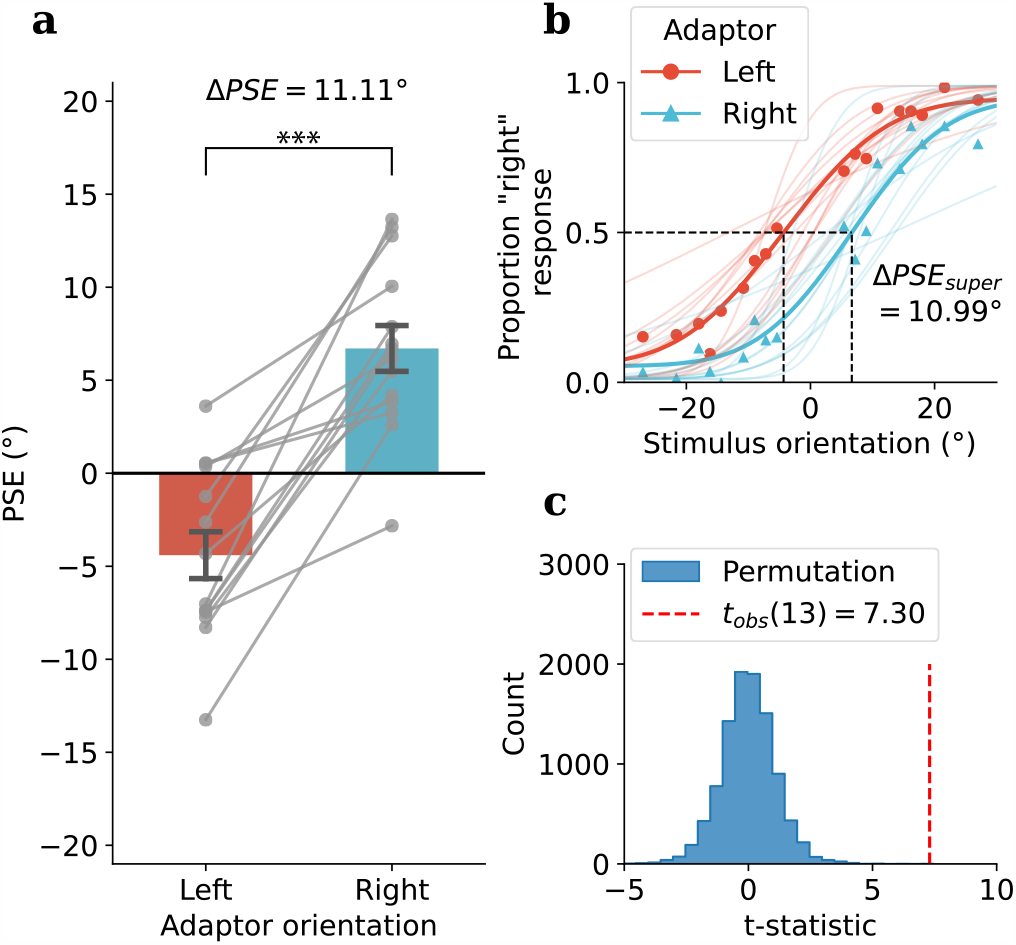
Experiment 1: Tactile TAE results: **(a)** Group average of PSEs following leftward (−27°, red bar) and rightward (27°, blue bar) tilted tactile adaptor, PSEs for individual participants are shown as grey dots. **(b)** Psychometric functions for each adaptor orientation (super-subject: bold lines, individual participants: faint lines). **(c)** A permutation test was used to assess the difference between PSEs under each adaptor orientation. The histogram of the t-statistics from 10,000 permutations of the PSEs is shown in blue, while the observed t-statistic is represented by the red dashed line, *p*_*perm*_ *<* .0001^***^.

### Crossmodal transfer of orientation adaptation from touch to vision

In the previous experiment, we successfully demonstrated a robust TAE in tactile orientation perception, similar to its visual counterpart. Given the close connection established by prior research in orientation perception between the two modalities, we aimed to investigate whether adaptation in one modality would induce TAE in the other modality.

Participants were tested in two conditions in Experiment 2: tactile adaptation, visual test (TV) and visual adaptation, tactile test (VT); In the TV condition, similar to Experiment 1, participants were first adapted using the titled tactile grating during the adaptation phase. In order for the participant to focus their attention on resolving the tactile orientations, which is thought to be critical in deploying the visual cortical process in tactile contexts^12^, participants were adapted through a series of pseudo-randomized two-interval-forced-choice (2IFC) tactile grating orientation discrimination trials with stimuli centred around the adaptor orientation (−27°or 27°), alternating between blocks and counter-balanced between subjects (see Method section for more details). After the adaptation period (initial: 60 s, top-up: 10 s, see Fig. 4), participants were then tested in vision using orientation-filtered pink noise stimuli (Fig. 1b). After each tactile adaptation phase, a series of six visual test trials were presented in the test phase. Psychometric functions were fitted separately for each trial (i.e., all first-trial responses, all second-trial responses, etc) for each participant. The PSEs from the first-trial data (immediately after adaptation) were used to assess the adaptation effect (as shown in Fig. 5), and the remaining five trials in the test phase were used to examine how the adaptation effect changed over time (Discussed in more detail in the last part of the Results section).

**Figure 4.**
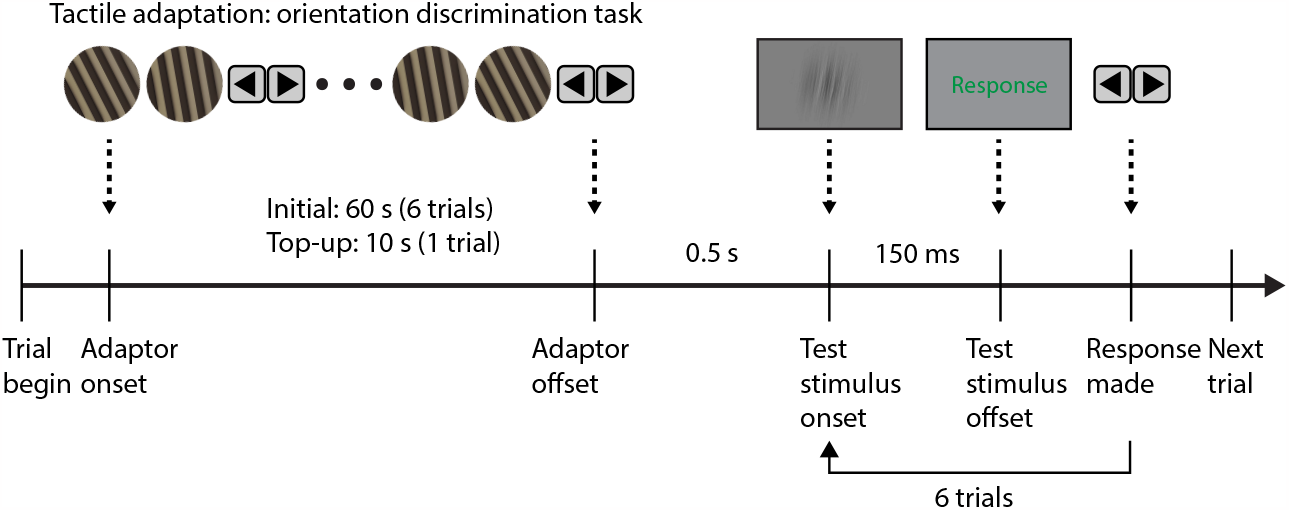
Procedure for Experiment 2, Condition 1: tactile adaptation, visual test (TV): during the adaptation phase (initial: 60 s, top-up: 10 s), participants were cued to perform multiple tactile orientation discrimination tasks, with the pseudo-randomised tactile stimulus that sits around either -27°(leftward adaptation condition) or 27°(rightward adaptation condition), after which participant went through the test phase that consists of a sequence of six visual test stimulus, with each stimulus presented for 150 ms.

**Figure 5.**
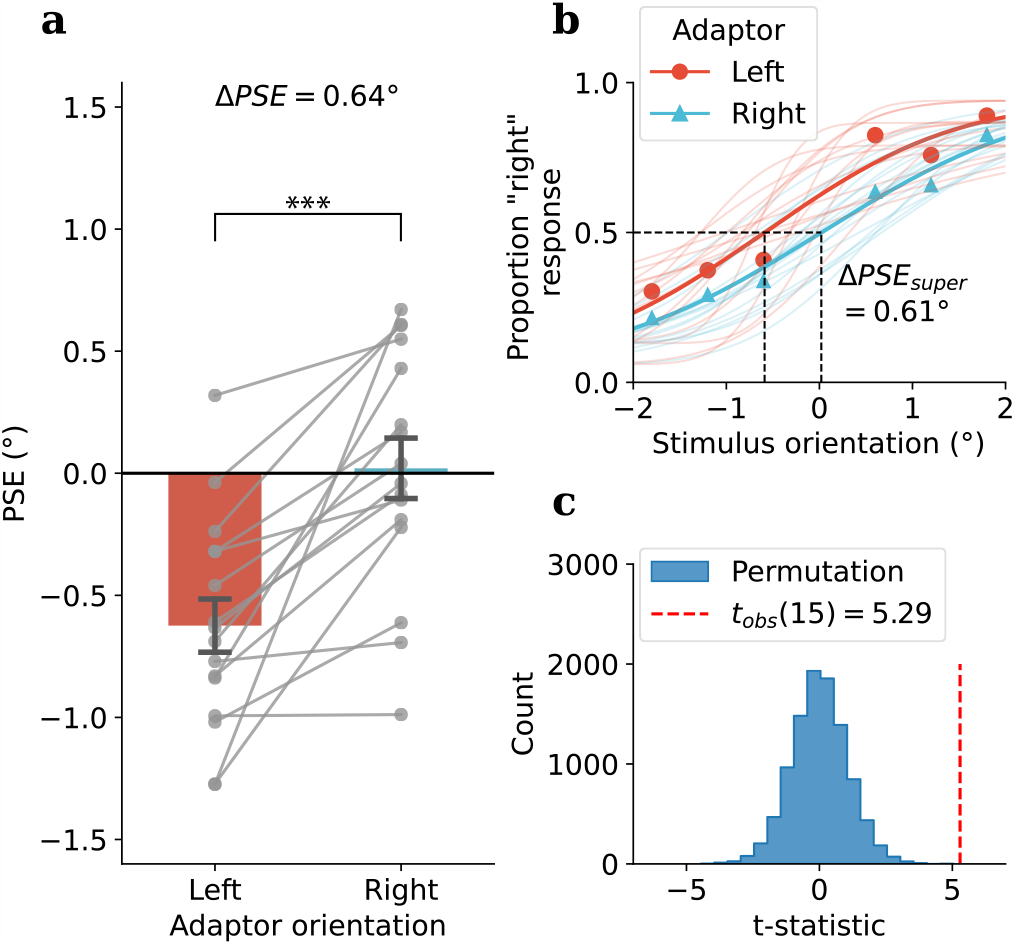
Experiment 2, Condition 1: tactile adaptation, visual test (TV) results: **(a)** Group average of PSEs in visual test following leftward (average -27°, red bar) and rightward (average 27°, blue bar) tilted tactile adaptor, individual participant PSEs were shown as grey dots. **(b)** Psychometric functions for each adaptor orientation (super-subject: bold lines, individual participants: faint lines). **(c)** A permutation test was used to assess the difference between PSEs under each adaptor orientation. The histogram of the t-statistics from 10,000 permutations of the PSEs is shown in blue, while the observed t-statistic is represented by the red dashed line, Holm-Bonferroni correction was applied to the permuted p-value, *p*_*Holm*;*perm*_ = 0.0006^***^.

Similar to Experiment 1, a significant difference in PSE was observed between leftward and rightward tactile adaptation conditions (Mean Δ*PSE* = .64,*SD* = .49, *t*_*obs*_(15) = 5.29, *p*_*obs*_ *<* .001, Cohen’s *d*_*obs*_ = 1.32, *p*_*holm*;*perm*_ = .0006^***^, see Fig. 5), which clearly shows that the TAE can transfer crossmodally from somatosensation to vision. Again, the repulsive effect was robust and present at the individual level for all 16 participants (Fig. 5a).

### Asymmetry in crossmodal tilt aftereffect: visual adaptation does not bias tactile orientation perception

In the second condition of Experiment 2, we tested the crossmodal transfer of orientation adaptation in the other direction — from vision to touch. Participants were adapted using either a leftward tilted (−27°) or rightward tilted (27°) orientation filtered pink noise pattern (see Fig. 1b for an example). A dot discrimination task on the stimulus (see Fig. 6) was employed in the adaptation phase to make sure the participant fixated on the visual adaptor. After the adaptation phase, participants were then tested in touch using the tactile grating. Each test phase consists of two trials, and the responses of the first trial were used to evaluate the crossmodal TAE here. And surprisingly, different from the previous condition, no significant difference in PSE was found between the two adaptor orientations (Mean Δ*PSE* = −1.34°, *SD* = 4.06°, *t*_*obs*_(15) = −1.32, *p*_*obs*_ = 0.20, Cohen’s *d*_*obs*_ = −0.33, *p*_*holm*;*perm*_ = 0.20, see Fig. 7).

**Figure 6.**
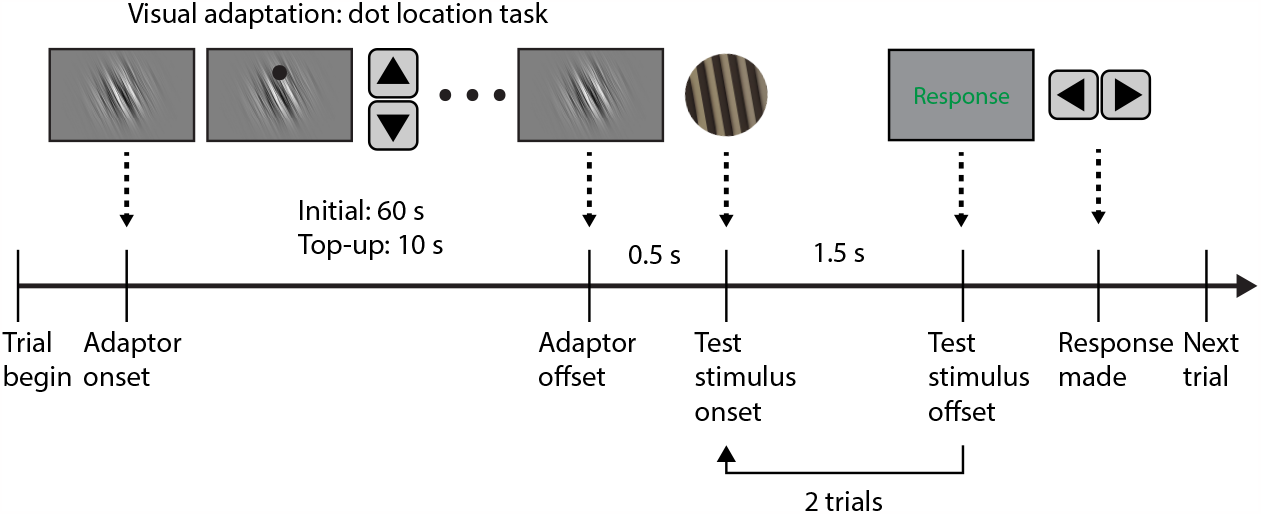
Procedure for Experiment 2, Condition 2: visual adaptation, tactile test (VT): during the adaptation phase (initial: 60 s, top-up: 10 s), participants were instructed to fixate on the orientation-filtered pink noise pattern (−27°or 27°), during which they were also instructed to perform a dot location task, to ensure there fixation on the visual adaptor. After this participant went through the test phase which consisted of a sequence of two tactile test stimuli, with each test stimulus presented for 1.5s.

**Figure 7.**
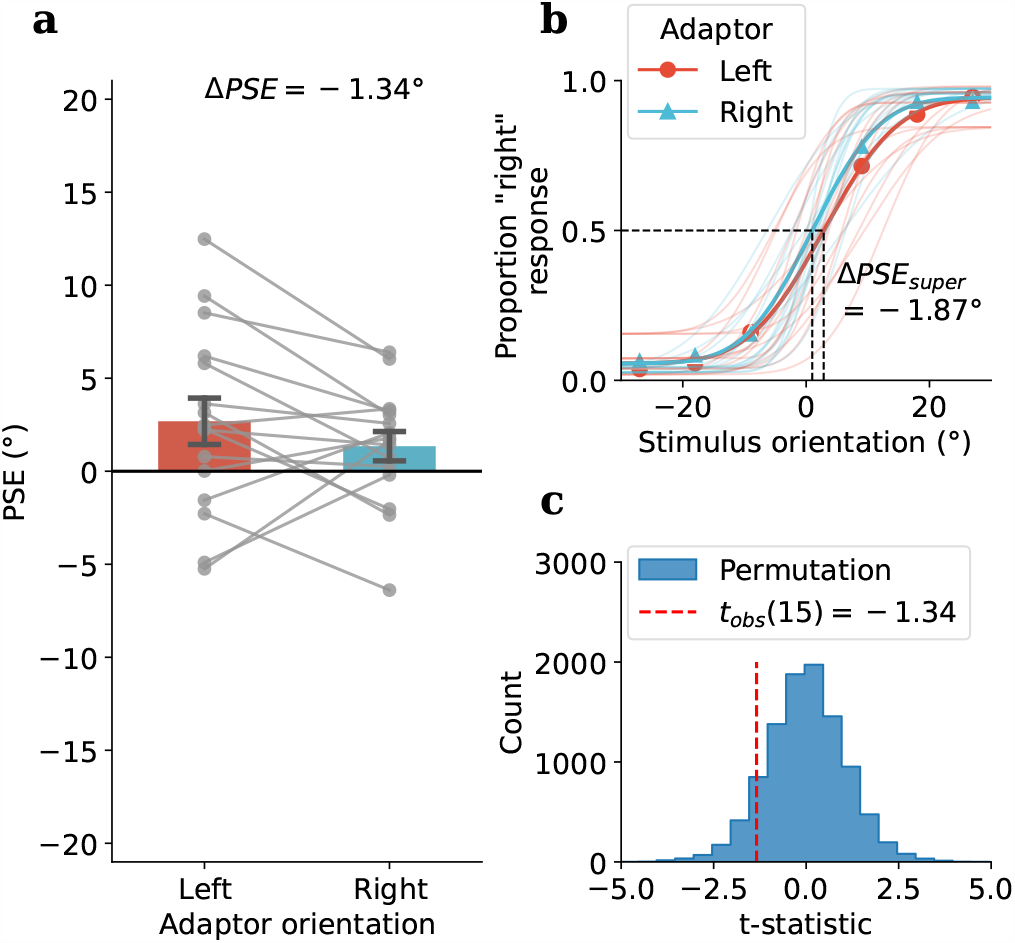
Experiment 2, Condition 2: visual adaptation, tactile test (VT) results: **(a)** Group average of PSEs in tactile test following leftward (−27°, red bar) and rightward (27°, blue bar) tilted visual adaptor, individual participant PSEs were shown as grey dots. **(b)** Psychometric functions for each adaptor orientation (super-subject: bold lines, individual participants: faint lines). **(c)** A permutation test was used to assess the difference between PSEs under each adaptor orientation. The histogram of the t-statistics from 10,000 permutations of the PSEs is shown in blue, while the observed t-statistic is represented by the red dashed line, Holm-Bonferroni correction was applied to the permuted p-value, *p*_*Holm*;*perm*_ = 0.20.

### Trial-by-trial variability in crossmodal tilt aftereffect and asymmetrical serial dependence

In Experiment 2, the adaptation phase had the same durations for both TV and VT conditions. However, a notable difference was that the test stimulus presentation time for tactile was 1.5 s — 10 times longer than that for visual test (150 ms). This difference in test duration can be attributed to the higher sensitivity of vision (mean orientation bandwidth *σ* = 1.29°, obtained from the fitted cumulative Gaussian psychometric functions) compared to somatosensory perception(*σ* = 7.84°). To account for the difference in stimulus presentation time, we decided to incorporate a sequence of test trials in the test phase following each adaptation phase. This approach allows us to examine the time course of changes in TAE after adaptation more comprehensively (See Fig. 4 & 6).

Similarly, psychometric functions were fitted for each of the remaining trials after adaptation, and permutation tests were performed to examine the TAE for each trial after adaptation, with p-values adjusted using the Holm-Bonferroni procedure. In condition 1 (TV), as stated earlier, a significant repulsive TAE was observed in the first visual test trial after tactile adaptation, however, this effect seems to diminish quickly after the first test trial, with no significant biases observed in the remaining trials (See Fig. 8a). In condition 2 (VT), no significant biases were observed for either the first test trial after adaptation or the second one (See Fig. 8b).

**Figure 8.**
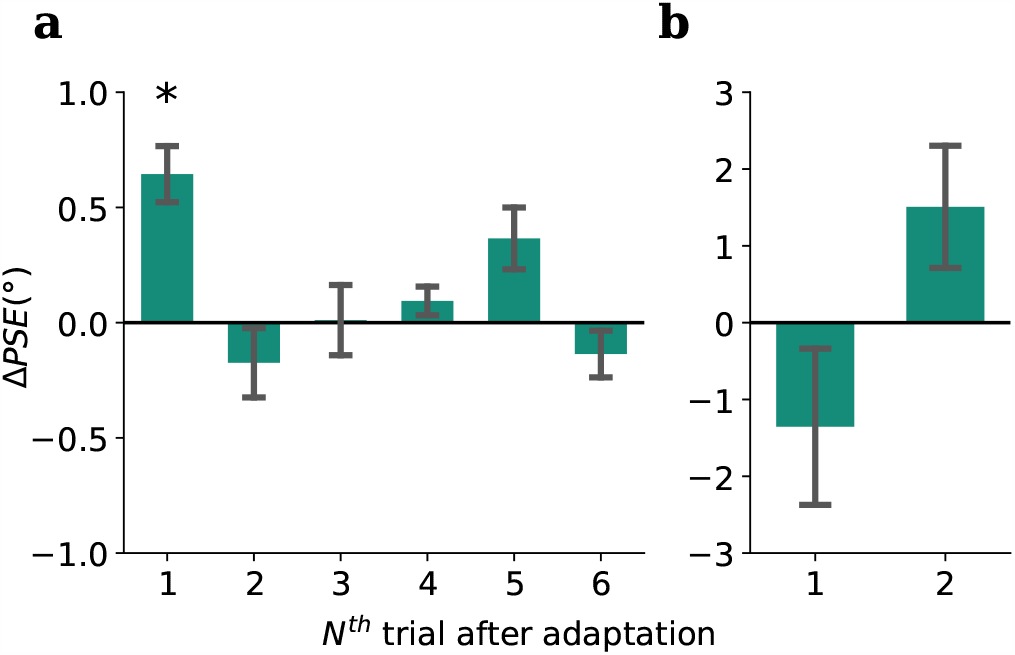
Experiment 2 results: **(a)** Condition 1, tactile adaptation, visual test (TV): after each adaptation phase, a sequence of six test trials were performed, the detailed analysis for the first trial immediately after adaptation was shown in Figure 5, similar permutation test were performed for all six trials, and the permuted p-values were corrected using Holm-Bonferroni procedure. A significant repulsive TAE were found for the 1st trial (*p*_*Holm*;*perm*_ = .0006^***^, see Fig. 5), while subsequent trials yielded non-significant p-values of *p*_*Holm*;*perm*_ = .58, *p*_*Holm*;*perm*_ = .95, *p*_*Holm*;*perm*_ = .73, *p*_*Holm*;*perm*_ = .07, *p*_*Holm*;*perm*_ = .58, respectively. **(b)** Condition 2, visual adaptation, tactile test (VT): no significant TAE were observed for either the 1st or the 2nd trial after adaptation, *p*_*Holm*;*perm*_ = .20, *p*_*Holm*;*perm*_ = .15 respectively.

In Condition 1 (TV), the absence of a TAE in the second and subsequent trials could simply be due to recovery from adaptation over time. However, it could also be a result of the adaptation effect being mediated by the percept of the previous test trial, a serial dependence^36–38^ effect might be operating in addition to the adaptation (especially given that we tested a sequence of test trials).

To evaluate the potential biases caused by serial dependence using PSEs, the trials need to be further divided based on 1-back stimulus orientation, which would lead to insufficient data points for proper psychometric function fitting, hence we instead used Generalized Linear Mixed Effect Model (GLMM) to evaluate the potential influence of serial dependence^39^. In Condition 1 (TV), the GLMM was fitted with data from the 2nd to 6th visual test trials from all participants. For Condition 2 (VT), GLMM was fitted for the 2nd tactile test trial. See the Methods section for details on the GLMM.

The results indicate that both the current stimulus and previous stimulus were significant predictors for the remaining test trials in Conditions 1 and 2 (see Table 1 for Condition 1 (TV) & Table 2 for Condition 2 (VT)). However, no significant difference in response was observed between the two adaptor orientations, for both TV and VT conditions. For graphical illustration purposes, we fitted a super-subject psychometric function for each combination of adaptor orientation and 1-back stimulus orientation direction (see Fig. 9). The significant serial dependence (difference in PSE between solid lines and dotted lines) and non-significant TAE (difference in PSE between blue and red lines) can also be clearly visualized in the figure. However, surprisingly, both the GLMM and the super-subject psychometric function have shown an asymmetry in the direction of the serial dependence between conditions, with TV showing repulsive serial dependence (*β* = −0.10, *t*(7196) = −7.90, *p <* .001^***^) and VT indicating attractive serial dependence (*β* = 0.01, *t*(2876) = 5.57, *p <* .001^***^).

**Table 1.** GLMM model statistic for remaining test trials (2nd to 6th following adaptation), in Condition 1 (TV) of Experiment 2.

| Name | Estimate | SE | t | df | p | 95% CIs |  |
| --- | --- | --- | --- | --- | --- | --- | --- |
|  |  |  |  |  |  | Lower | Upper |
| (Intercept) | 0.106 | 0.053 | 2.000 | 7196 | 0.046* | 0.002 | 0.210 |
| Current Stimulus | 0.538 | 0.044 | 12.314 | 7196 | < 0.001*** | 0.453 | 0.624 |
| Previous Stimulus | -0.100 | 0.013 | -7.895 | 7196 | < 0.001*** | -0.124 | -0.075 |
| Adaptor Orientation | -0.023 | 0.033 | -0.714 | 7196 | 0.476 | -0.087 | 0.041 |

**Table 2.**
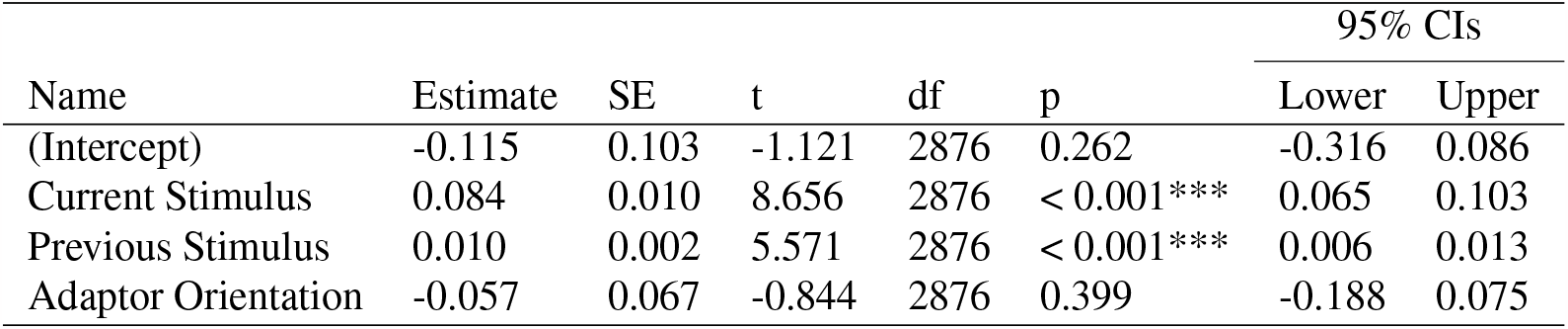
GLMM model statistic for the second test trial (following adaptation), in Condition 2 (VT) of Experiment 2.

| Name | Estimate | SE | t | df | p | 95% CIs |  |
| --- | --- | --- | --- | --- | --- | --- | --- |
|  |  |  |  |  |  | Lower | Upper |
| (Intercept) | -0.115 | 0.103 | -1.121 | 2876 | 0.262 | -0.316 | 0.086 |
| Current Stimulus | 0.084 | 0.010 | 8.656 | 2876 | < 0.001*** | 0.065 | 0.103 |
| Previous Stimulus | 0.010 | 0.002 | 5.571 | 2876 | < 0.001*** | 0.006 | 0.013 |
| Adaptor Orientation | -0.057 | 0.067 | -0.844 | 2876 | 0.399 | -0.188 | 0.075 |

**Figure 9.**
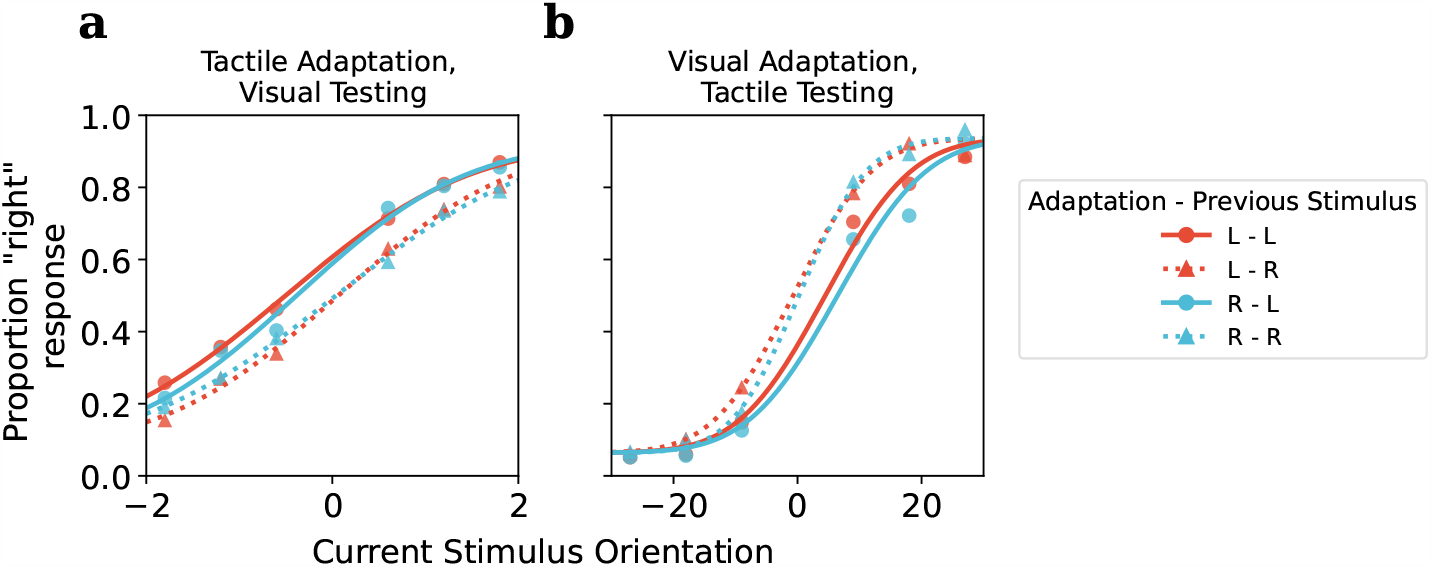
Super-subject psychometric functions by adaptor orientation (denoted by colour) and previous stimulus orientation (denoted by solid and dotted lines). Previous stimuli were grouped based on their direction for this plot (left vs. right), while the actual orientation in degree is used in the GLMM model. The purpose of this plot was to graphically illustrate the findings of the GLMM. **(a)** Condition 1: tactile adaptation, visual test, a repulsive serial dependence (difference between solid & dotted lines), but no TAE were observed (between red and blue lines), which is confirmed by GLMM results. **(b)** Condition 2: visual adaptation, tactile test, an attractive serial dependence (difference between solid & dotted lines), but no TAE were observed (no difference between red and blue lines), which is also confirmed by GLMM results.

## Discussion

In this study, we used the classic tilt aftereffect paradigm to test adaptation to orientation in the somatosensory system. Our first goal was to determine whether a TAE would occur in the touch domain (Experiment 1) and whether it would exhibit the repulsive tuning that is observed in the visual TAE. Our second goal, motivated by accumulating cortical and behavioural evidence for visual-somatosensory interactions^7–15^, was whether any adaptation to orientation would transfer crossmodally between vision and touch (Experiment 2). In Experiment 1 we successfully demonstrated a robust tactile TAE using a tactile grating which displayed the typical repulsive effect observed with the visual TAE. The tactile TAE was robust and observable reliably at the level of the individual in all participants (Fig. 3). Experiment 2 tested whether there is a crossmodal transfer of adaptation effect between the two modalities, and revealed an interesting asymmetry in which tactile adaptation to orientation induced a repulsive TAE when tested in vision, yet adaptation to visual orientation did not generate a tactile TAE. We also examined the time course of crossmodal adaptation and found the repulsive tactile-to-vision TAE diminished quickly and was no longer significant after the first trial in the visual test sequence. Finally, we examined whether there were serial dependence effects in the test sequences used in the crossmodal conditions (Experiment 2). In another curious asymmetry, serial dependence was found in both VT and TV conditions, however, it was a repulsive effect for visual test sequence and an attractive effect for tactile test sequence.

The significant tactile TAE found in Experiment 1 is in agreement with a very recent report using a different paradigm to study tactile orientation. Hidaka et al.^33^ presented orientation on the palm of the hand using a two-point stimulation task. The alignment of two points was sufficient to produce a percept of orientation and adaptation to this stimulus produced a robust TAE. Somewhat surprisingly, these are the first two published studies to document a purely tactile TAE (i.e., adapting and testing with tactile stimuli). While our study is focused primarily on crossmodal transfer of adaptation, Hidaka et al. made important observations relating to somatosensory processing of orientation by showing that the TAE did not depend on hand rotation and that adaptation transferred from the palm to the back of the hand (although not between hands), suggesting it is not coded in an external reference frame. Our results complement Hidaka et al.’s report by adding that orientation on the fingertip also produces a TAE that is very robust and apparent at the individual level in all participants. Importantly for our purposes, the robust tactile TAE we observed confirms the feasibility of using the same tactile stimulus settings to explore crossmodal TAEs between touch and vision in Experiment 2.

Our second experiment examined crossmodal transfer of orientation adaptation between touch and vision, testing for both vision-to-touch and touch-to-vision transfer. As shown in Figure 5, there was a very clear effect of tactile adaptation to orientation on the fingertip transferring to vision and causing orientation repulsion on visual test stimuli. The effect was very reliable and was present at the individual level in all participants. This result is consistent with the literature reviewed in the Introduction showing behavioural evidence of common processing mechanisms for visual and tactile orientation. For example, a tactile grating combined with a visual surround grating increases the magnitude of the tilt illusion^35^, showing a clear orientation interaction, and visual-tactile orientation interactions have been shown to exhibit surprisingly tight orientation tunings^21,22^, possibly due to the increased perceptual precision that arises from integration of multisensory cues^40^. Despite these behavioural findings, and a host of neural findings supporting visual-tactile interactions^12,13,16^, the crossmodal transfer of adaptation did not occur in the reverse direction. As shown in Figure 7, there is no evidence of transfer of visual orientation adaptation to tactile orientation perception. There is no definitive explanation for this null result (which we consistently observed during several pilot experiments testing various parameters), although we consider several possibilities below.

### Potential explanation for asymmetry in crossmodal TAE and implications for crossmodal connections

As a first observation, it is clear that orientation signals from the two modalities in our study were not simply combined in the manner of Bayesian maximum likelihood estimation (MLE) integration. MLE fusion of multisensory signals does occur and is well documented in the perception of audio-visual spatial location^41^ and visual-tactile size^42^, but it does not appear to happen for visual-tactile orientation. The reason for this may relate to the very different orientation sensitivities in touch and vision, with orientation discrimination in touch being much poorer than in vision. Behavioural studies show that human orientation discrimination in vision is typically well below 1°for a wide range of spatial frequencies and can be as low as 0.3°– 0.5°in the most sensitive range^43–45^. This is around 10 times better than human orientation discrimination in touch where the best-reported estimates are 4.2°– 5.4°for static and passively applied stimuli on the finger pad^46^. Our behavioural observations are consistent with these neural findings in that we found a mean bandwidth for visual orientation of 1.29°(average value of *σ* from the fitted psychometric functions) and a mean bandwidth for tactile orientation of 7.84°. These perceptual results reflect single-unit neurophysiological data recorded from primates. In the primary visual cortex, most neurons are orientation-tuned and the full bandwidth is typically in the range of 30°-40^°47^. In the somatosensory cortex, neurons selective for orientation on individual finger pads are found in S1 and across multiple fingers in S2^6^. An example of an S1 neuron with a “sharp tuning” is shown in Hsiao et al^6^ and has a full bandwidth of about 50°. In the second somatosensory cortical area (S2) the average orientation tuning bandwidth is 63^°48^. Given these marked differences in orientation precision, an MLE framework would always predict that vision would dominate visual-tactile orientation perception. Thus, in the TV condition (where we found a strong transfer of tactile orientation adaptation to vision), a Bayesian fusion model would predict little change in post-adaptation orientation as vision would be by far the higher weighted input. Thus, vision would be expected to dominate orientation perception regardless of the adapted state of tactile orientation neurons, yet against this, we found a strong TAE in the TV condition. Conversely, the MLE fusion model would predict strong a TAE in the VT condition, as the adapting stimulus was visual and thus in a vision-dominated fusion model the adapted state of visual neurons should be clearly evident, yet we did not observe a TAE.

Another consideration is that there may have been a genuine TAE in the VT condition but it was simply missed. Neural and methodological factors could both contribute to this possibility. Because of their markedly different orientation bandwidths, finding evidence of orientation repulsion following visual adaptation (which typically produces a visual TAE on the order of 1°– 2.5^°29,49^ in the tactile system where the level of orientation precision is much poorer might simply lead to a small effect transferred from vision to touch being too noisy to measure reliably. This is compounded by the fact that aftereffects are relatively brief and decay over time, returning continuously towards their unadapted baseline. This makes it difficult methodologically to measure the tactile aftereffect as measurements in touch generally take more time than in vision. In vision, it is easy to flash a brief test grating and quickly probe the adapted state of the orientation system but in touch, this is usually a much slower process. In our experiment, the tactile grating was presented for a duration of 1.5 s in the test condition and it is possible there may have been a transfer of orientation adaptation from vision to touch in the VT condition but it dissipated and fell below the threshold during test trials as participants felt the stimulus and arrived at their decision about its orientation. If the decay of the aftereffect was a critical factor in our failure to observe the transfer of orientation adaptation from vision to touch, it remains possible that it could be obtained under different test conditions using briefer tactile test trials. Supporting the possibility that we simply missed a genuine TAE in the VT transfer condition is the fact that this same condition has been successfully demonstrated by Krystallidou and Thompson^34^, although they tested tactile orientation on the forehead and used a two-point test stimulus. It remains to be seen whether the same effect would be obtained when testing as we did, on the finger pad with oriented gratings and we propose that brief tactile test stimuli would likely improve the probability of detecting the effects of visual orientation adaptation in touch.

By contrast, the transfer of orientation adaptation from touch to vision was robust. How this would arise neurally is not well established. One possibility is that tactile-visual interactions for orientation are mediated by imagined or visualized orientation when engaged in touching tactile orientation, a suggestion made by Sathian and Zangaladze^12^. Indeed, multiple studies have shown visual imagery is capable of inducing aftereffects through top-down recruitment of the visual cortex^50–52^. Particularly, Mohr et al^53,54^ showed that mental imagery alone was sufficient to produce orientation-specific adaptation in the extrastriate visual cortex (V3-V4) and elicit a consequent visual TAE. Thus if tactile orientation were to interact with vision through visualizing the felt orientation^12^, it would still provide a possible means for the transfer of orientation adaptation from touch to vision that we observed in the visual test condition.

Interestingly, Ganis and Schendan showed that mental imagery and perception produce opposite-signed adaptation effects in face perception^50^ (with the imagery effect being attractive). Mohr et al. studied TAEs with imagined visual stimuli and found a similar result: while imagined lines produced the classic direct TAE(repulsion of orientations near the adapted orientation), they found the indirect TAE(where orientations far from adaptation are tested) had the opposite sign to the standard attractive effect elicited directly from visual adaptation^53^. This also seems to coincide with our results in the visual-to-tactile condition, which though not significant, did point to an attractive TAE while testing in touch (Fig. 7), although further work would be needed to establish this concretely. It seems that the top-down recruitment of the visual cortex might lead to the opposite adaptation effects (attraction), but the cause of this is not clear yet, possibly it has to do with the differences in neural mechanisms between bottom-up and top-down recruitment of the primary visual cortex.

Another possibility is that the adaptation effect transfers via direct connections between primary cortices for touch and vision. This notion is appealing and sits well with recent challenges to the traditional view that the primary cortices are strictly unimodal and well segregated from each other^55,56^. Direct interactions between primary sensory cortices have been well documented in rodents^57–59^, however corresponding studies in primates show that direct projections between primary cortices are quite sparse^60–64^ and primates probably rely more on multisensory interactions in areas beyond primary cortices. This view squares with evidence for tactile input to the extrastriate visual cortex during tactile orientation discrimination such as Sathian et al.’s^16^ PET study and Zhang et al.’s^15^ fMRI study showing orientation discrimination elicited significant task-specific activations in the extrastriate visual cortex, and a TMS study by Zangaladze et al.^13^ showing the same area was critical to discriminating tactile grating orientation.

### Adaptation and serial dependence acting as competing mechanisms

In Experiment 2, the GLMM analysis on the remaining test trials (excluding the initial trial) reveals contrasting intramodal serial dependence effects in the two conditions. These serial effects were present while no crossmodal TAE was found in these remaining trials by the GLMM modal. Serial dependence has been widely studied in vision (for review, see Pascucci et al.^65^ and Cicchini et al.^66^), with orientation being one of the first visual features that have been looked at^36^. In condition VT, we were able to show an attractive intramodal serial dependence in the tactile orientation task, which is first up to date, and also seems to coincide with our hypothesis that the tactile orientation judgment shares a very similar processing mechanism as vision. In Condition TV, we did not observe a classical attractive serial dependence, on the contrary, we found a repulsive intramodal serial effect, which is very interesting.

Various previous studies looking at serial dependence also find a mixture of attractive and repulsive effects^67–73^, and suggests that this mixture of effect direction might be the result of two competing mechanism: lower-level repulsive sensory adaptation effect which was presented right after the previous percept, and attractive serial dependence that manifests throughout higher order post-perceptual processing^65^. The observed direction of the perceptual history effect at a certain time point was thought to be the result of the combination of these two competing mechanisms^65,73^. Previous studies also indicated that the strength of attractive serial dependence is dependent on the reliability of the current percept, in particular, the attractive effect is greater when the current precept lacks reliability (hence more weight is put on the recent perceptual history to optimize the current percept^37^). And in our experiment, orientation perception is much more reliable in vision (*σ* = 1.29°) compared to touch (*σ* = 7.84°). In condition TV, with the more reliable visual orientation judgment, the weight on the previous percept should be lower, which should lead to a weaker attractive serial effect. Besides, the immediate response cue after the brief presentation of visual test stimuli (150 ms) might be too early for the attractive serial dependence to manifest in the higher-order post-perceptual processing, which together could lead to the observed classical sensory-driven repulsive effect. It would also be able to explain the quick diminishing of crossmodal TAE in Condition TV: the visual neurons that were crossmodally adapted by tactile adaptor could be re-adapted intramodally by the visual test stimuli, and hence diminishing the touch-to-vision crossmodal TAE in the subsequent test trials.

In condition VT, tactile orientation judgment would potentially rely more on the previous percept due to the lack of reliability, and the much longer presentation time (1.5 s) would provide more time for higher-order post-perceptual processing, hence showing a stronger attractive intramodal serial effect at the time of cued response. The absence of vision-to-touch crossmodal TAE in this condition can also be attributed to this lack of reliability in tactile orientation judgment, as the effect of the weak repulsive crossmodal sensory adaptation being overpowered by the stronger attractive influence of perceptual history due to high uncertainty in perception, and fits with our suggestion that the hint of an attractive effect in our vision-to-touch TAE condition (Fig.7), and provides a good theoretical framework to motivate a reinvestigation of this condition in future studies to clarify whether there is a genuine attractive effect or not.

### Divisive normalization between modalities leads to a combination of repulsive and attractive effects

Figure 7 shows a mix of attractive and repulsive biases among participants for condition VT. These effects might simply reflect noise in a genuine null condition, but they might also indicate a genuine mixture of adaptation-induced attractive and repulsive biases. Some studies in vision have reported attractive adaptation shifts with certain paradigms^74,75^, although repulsive effects are more common. One flexible model used to explain adaptation effects is the divisive normalization model and it can explain the difference in the direction of adaptation-induced bias across many different sensory adaptation paradigms^75–77^. At the neuronal level, divisive normalization is proposed to work by taking a given neuron’s activation from within its own receptive field (Classical Receptive Field - CRF) and dividing it (i.e., normalising it) by the sum of activation from neurons arranged in a larger suppressive surround field (normalization pool). The net effect is that stimulation in the normalization field down-regulates the neuron’s response. Phenomena such as surround suppression observed in V1 can be explained by divisive normalization as extending the stimulus extent into the surround field reduces a neuron’s response^76^. Divisive normalization has been used to model many perceptual and neurophysiological effects and is also thought to play a role in multisensory integration^78^. Hence it may play a role in how orientation information is transferred from somatosensory to the visual cortex. We tried to simulate a divisive normalization model based on these principles where an orientation-selective visual neuron has a normalization field with a broader orientation range compared to the CRF. We were able to show that the sign of the adaptation effect – whether attractive or repulsive – could vary based on the relative strength of the adaptation drive to the CRF and normalization field (See Fig 10). In condition VT, the adaptation drive to the CRF and normalization pool of the visual neurons that was later involved in the tactile test might differentiate between subjects, and hence lead to the mixed directions in the bias. However, the model and simulation here were only for illustrative purposes to demonstrate the possible explanation of the observed behaviour, future research with actual neuronal recording might help confirm this hypothesis.

**Figure 10.**
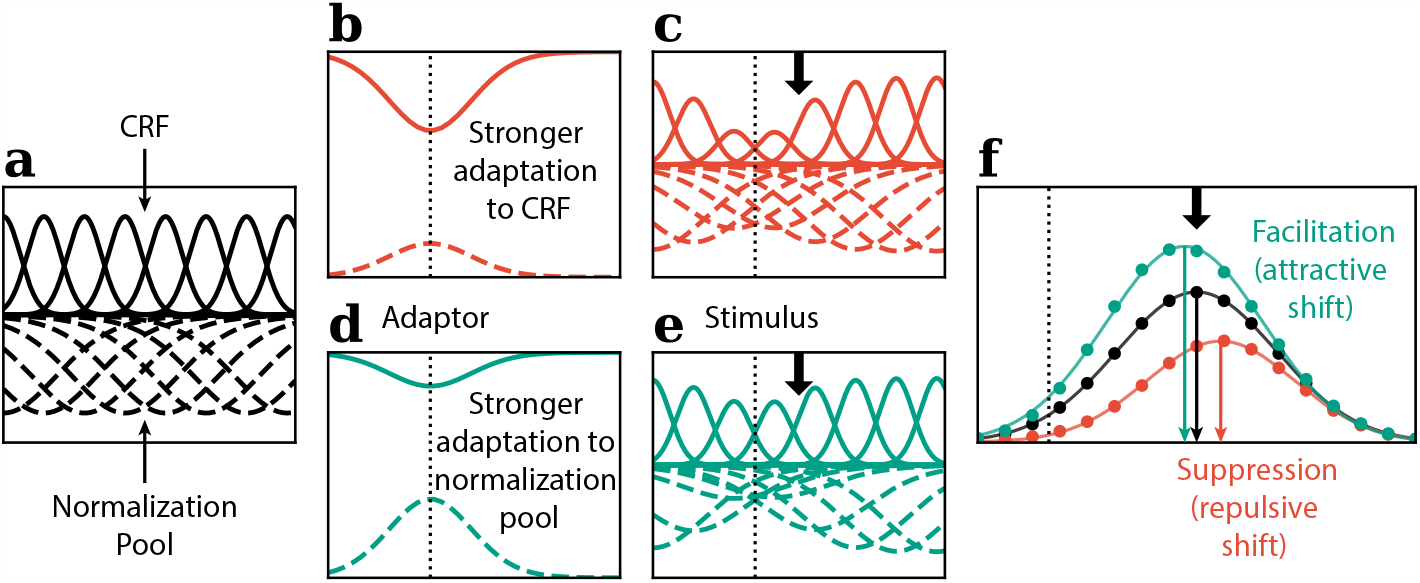
Illustration of the Divisive normalization model.**(a)** CRF and normalization pool prior to adaptation. **(b-c)** Stronger adaptation to the CRF **(d-e)** stronger adaptation to the normalization signal **(f)** Stronger adaptation to the CRF lead to classic repulsive TAE with suppression effect on the population response, while stronger drive to the normalization signal lead to attractive adaptation and facilitating effect in the population response, which could potentially explain the asymmetrical crossmodal TAE results in our experiment.

## Conclusion

In the current study, we found that adaptation to a tactile grating presented to the finger pads produced a robust, repulsive TAE, analogous to what is observed in vision after adaptation to a visual grating. We also explored the transfer of orientation adaptation between vision and touch. Adaptation to tactile orientation produced a very robust TAE when tested in vision that was reliable at the level of individual participants. This result is consistent with mounting neural evidence of common processing of visual and tactile orientation in the extrastriate cortex. However, in an asymmetrical result, we found no evidence of transfer of orientation adaptation in the other direction, from vision to touch. We propose that the failure to find an effect of visual orientation adaptation when test in touch is inconclusive and should be interpreted with caution. It is inconsistent with a behavioural report of vision-to-touch transfer of orientation adaptation (albeit for test stimuli presented on the forehead rather than the finger pad). We conjecture that our rather long tactile test trials may have prevented effective measurement of transferred orientation adaptation and recommend future studies employ brief test trials to better capture the ephemeral and decaying nature of orientation adaptation. It is also possible that the poor precision of tactile orientation perception relative to vision adds to the difficulty of capturing vision-to-touch transfer of orientation adaptation. We also looked at the intramodal serial dependence effect within the test sequence and found a similar asymmetrical serial dependence effect. Which could be a competing mechanism that mediates the crossmodal TAE, and provides another alternative explanation for the asymmetrical crossmodal transfer of TAE. Finally, we propose that a divisive normalization model could explain our inconclusive findings in the case of vision-to-touch transfer as both attractive and repulsive shifts can be induced by the adaptor depending on the nature of the surround field. Thus our null result might reflect roughly equivalent numbers of subjects showing a repulsive or an attractive effect.

Despite these caveats, we have demonstrated two key findings: (i) for the first time we have shown a purely tactile TAE on the finger pads, and (ii) we show clear evidence of crossmodal transfer of adaptation from touch to vision, complementing an earlier report of transfer from vision to touch. The first result complements neural work in primates showing orientation selectivity in S1 neurons and indicates that neural interactions among S1 cells with adjacent orientation preferences interact similarly to those observed in the visual cortex. The second result adds to mounting neural and behavioural evidence of common processing of visual and tactile orientation.

## Methods

### Participants

14 naïve participants participated in Experiment 1. 22 naïve participants participated in Experiment 2. All participants reported being right-hand dominant with no recent history of damage to the right index finger and no history of damage or diseases of the nervous system.

### Apparatus & stimulus

The tactile stimulus used in all experiments was a 3D printed (Ultimaker 3D printer) disk with a grating pattern on its upper surface whose orientation could be rotated with an Arduino Uno microcontroller. The model of the grating disk was designed using FreeCAD. It has a diameter of 30 mm and is composed of a triangular wave with a 6 mm spatial period and a peak-to-trough amplitude of 2.5 mm (See Fig. 1c). The microcontroller is controlled by an Arduino Uno Board via MATLAB and can rotate the disk with a resolution of 1.8°/step. The disk was housed in a 3D printed box on which the subject rested their hand. The disk was presented to the subject through an aperture on top of the box and the presentation period was controlled by raising the disk into position for a fixed period and then lowering it back inside.

Visual stimuli were generated using a MATLAB image processing toolbox^79^, and presented using Psychophysics Toolbox extensions^80^ on a Dell OptiPlex 7440 Display (1920 *×* 1080 pixels, 60 Hz refresh rate). Visual adaptation and test stimuli in these experiments1 were orientation-filtered noise patterns (pink noise, filtered with a Gaussian orientation filter with bandwidth (*σ* = 5°), with a 2D Gaussian kernel applied to the noise pattern to generate a smooth roll-off at the edge. The root-mean-squared (RMS) contrast of the gratings for the visual adaptor used in the VT condition was 0.3, and for the test stimuli in the TV condition it was 0.018.

### Data analysis

#### Exclusion Criteria

For Experiment 1, participants with an exceptionally low percentage of correct performance(*<* 65% in the 2AFC task were excluded from the analysis, as reliable psychometric functions could not be fitted to assess the biases in PSEs. For Experiment 2, similar exclusion criteria were applied, participants with performance lower than 65% correct in either of the 2 conditions were excluded from the analysis.

#### Psychometric function fitting

Psychometric functions were fitted for each participant and each adaptor orientation separately:

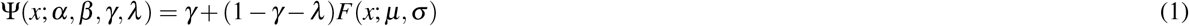

The function *F*(*x*; *μ, σ*) is a cumulative Gaussian function where *μ* =mean and *σ* =standard deviation. And *γ & λ* represent the guess rate and the lapse rate. These parameters correspond to the disparity between the asymptotes of the psychometric function and the ideal performance levels of 0% and 100%, respectively. Point of subjective equality (PSE) was obtained as the mean of the cumulative Gaussian function(*μ*, 50% rightward response), which was used as a measure for bias in orientation judgement (positive: leftward bias, negative: rightward bias).

#### Permutation Test

We used a non-parametric permutation test to evaluate the biases in the PSEs. The observed statistics (*t*_*obs*_) were obtained through paired-sample t-tests comparing the PSEs of the actual participants’ data under different adaptor orientations. Then, 10,000 permutations of the empirical PSEs were generated by randomly swapping the PSEs under different adaptor conditions. The p-value of the permutation test is calculated as the proportion of t statistics from the permuted distribution that is more extreme than the observed statistics(| *t*_*permutation*_| ≥ | *t*_*obs*_|).

For each condition, as multiple trials were tested after the adaptation, and similar permutation tests were performed for each test trial, we used the Holm-Bonferroni method^81^ to correct for family-wise error rate. The adjusted p-value is calculated by sorting the permuted p-value in each condition in ascending order, and multiplying the p-value with the number of tests (*C*) in the condition minus the rank(*i*) and plus one, the formula is as follows:

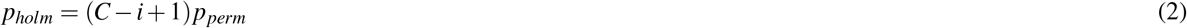

where an adjusted p-value larger than one is set to one.

#### Generalized Linear Mixed Effects Model (GLMM)

The serial dependence effects were analysed using GLMM, as further dividing each participant’s data based on previous stimulus in addition to adaptor orientations would result in insufficient data for proper psychometric function fitting, and lead to unreliable PSE estimates.

The model was fitted with the raw trial-by-trial data, following the principle of Moscatelli et al.^39^ The first test trial after each adaptation phase was excluded in this analysis as there is no 1-back test trial (the previous test trial was prior to the preceding adaptation phase, which is separated with the test trial by 60 s/10 s for initial and top-up adaptation, and would not be appropriate for serial dependence analysis).

We employed MATLAB’s Statistics and Machine Learning Toolbox to fit the GLMM. The model was fitted using a binomial distribution and probit link function, applying the maximum likelihood method with Laplace approximation. We included the three fixed effects, to account for the previous and current stimulus, as well as the adaptor orientation. We also included the subject as a random effect to account for the differences in sensitivity and overall internal biases for each participant. The GLMM model is presented in Wilkinson notation below:

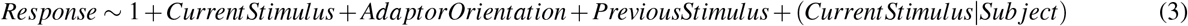

Notice the previous stimulus orientations used in the actual GLMM were in degrees. However, for clarity, in the graphical illustration (Fig. 9) the previous stimulus was simplified to left or right.

### Design & procedure

#### Experiment 1: tactile tilt aftereffect

Participants sat in front of a screen with their right hand resting on top of a box containing the tactile device. A schematic plot is shown in Figure 1a. Each session began with a preliminary test lasting approximately five minutes to determine for each participant which of the three orientation ranges was best suited to their tactile orientation acuity to ensure a better-fitting psychometric function (Fig. 1d). This also acted as the practice block to familiarize the participant with the task. The main test contains six blocks, with adaptor orientations (left vs. right) alternating between blocks, and the order counterbalanced between subjects. Each trial began with the tactile grating rising from inside the box to the level of the participant’s fingertip already rotated to the adaptor orientation used for that block. Participants passively felt the orientation for the duration of the adaptation phase (30 s for initial adaptation, 5 s top-ups thereafter). After the adaptation phase, and a gap of 500 ms, the test stimulus was then presented to the participant through the tactile device for 1.5 s, and the participant was cued to passively feel the orientation of the test stimuli with their fingertip, and then report whether the stimulus orientation was left (anticlockwise) or right (clockwise) compared to vertical (proximal-distal orientation) by pressing the arrow keys with the other hand (See Fig.2).

#### Experiment 2: visuotactile crossmodal tilt aftereffect

Experiment 2 consisted of two consecutive sessions, each testing a different condition: tactile adaptation, visual test (TV) and visual adaptation, tactile test (VT). The order of the two conditions was counterbalanced between participants. Both conditions consisted of an adaptation phase followed by a test phase.

In Condition TV, in order for the participant to focus their attention on resolving the tactile orientations during the adaptation phase, which is thought to be critical in deploying the visual cortical process in tactile contexts^12^. A 2IFC orientation discrimination task was introduced as the adaptation task: two tactile orientations were presented in succession, with each presentation lasting five seconds. Participants responded on the keyboard whether the second orientation was more towards the left or right of the first. All adaptation stimuli were pseudo-randomly drawn from four orientations (18°, 23.4°, 30.6°, 36°, left or right based on the block). At the beginning of each block, a sequence of six adaptation trials was performed, adding up to 60 seconds of initial adaptation. After each visual test sequence, another top-up adaptation trial was conducted, which accounted for 10 seconds of top-up adaption (See Fig. 4) For each block, the average of the pseudo-randomised adaptation trial stimulus orientation was -27°or 27°, alternating between blocks and counterbalanced between subjects.

In the VT condition, the visual adaptation phase contained a dot detection task designed to keep the participant’s attention on the adaptation stimulus. The dot appeared randomly in an upper or lower location multiple times during the adaptation period and the participant responded to the dot location with the keyboard every time they saw the dot, as shown in Figure6.

In both conditions, a test trial was presented immediately after the adaptation phase to test for the TAE. However, to accommodate the different nature of visual and tactile perception and the longer time needed to determine the tactile orientation, the test stimuli were presented for different durations in each condition (visual test: 150 ms, tactile test: 1.5 s). To evaluate the potential confound introduced by different time courses of testing, additional test trials were given after the first trial to test the change in TAE after the initial test. In Condition TV, five addition test trials were conducted (six test trials in total, See Fig. 4)), and in Condition VT, one addition test trial was conducted (two test trials in total, See Fig. 6).

### Divisive Normalization Model

To try and explain the mixed results in the visual adaptation, tactile test condition (VT), we used a simple divisive normalization model to simulate the population response of neurons in the visual cortex. This simplified illustrative model was inspired by some published data^75–77^.

The CRF (summation field) and normalization field (suppression field) tuning of each of the orientation-selective neurons is simulated with Von Mises functions (see Fig. 10a):

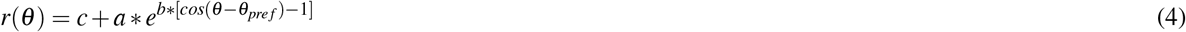

where c is an offset representing the base response, a is the peak height of the summation field/normalization field, and b is a parameter that controls the bandwidth of the summation field/normalization field. *θ*_*pre f*_ stands for the preferred orientation of the neuron.

The adaptation effect *a*(*θ*_*pre f*_) to a neuron with preferred orientation *θ*_*pre f*_ was simulated with the complement of the Von Mises function, with different strength (*k*) applied to the summation field and normalization field, and the strongest drive were observed at the adaptor orientation *θ*_*adaptation*_. The adaptation effects were then multiplied with the summation field and suppression field to obtain the tuning after adaptation (see Fig. 10b & d):

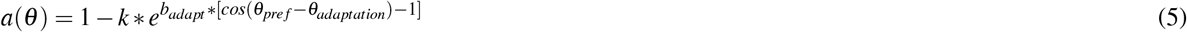

The normalized response *r* _*j*_ of the neuron *j* was calculated by dividing the CRF response (weighted sum of drive to the summation field ∑_*k*_ *ω*_*ik*_*I*_*k*_, where *I*_*k*_ was the non-normalized response of the neuron *k* in the summation field), divided by the normalization signal (weighted sum of drive to the normalization field 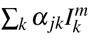). Constant *σ* was added to the denominator to avoid division by zero. *n & p* are exponents to account for non-linearity:

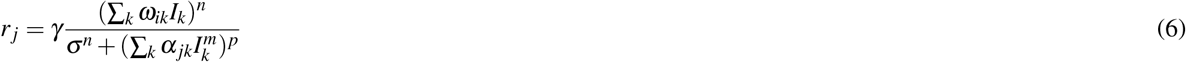

The population response was obtained by fitting a Von Mises function to the normalized responses of all the simulated neurons with different preferred orientations. The centre of the Von Mises function, representing the peak response, was used as the simulated perceived orientation (see Fig. 10f).

## Author contributions statement

D.A. conceived the experiments, G.W. coded and conducted the experiments, G.W. analyzed the results. All authors reviewed the manuscript.

## Data availability statement

The data that support the findings of this study are available on request from the corresponding author, G.W.

